# A 3-component module maintains sepal flatness in Arabidopsis

**DOI:** 10.1101/2023.12.06.570430

**Authors:** Shouling Xu, Xi He, Duy-Chi Trinh, Xinyu Zhang, Xiaojiang Wu, Dengying Qiu, Ming Zhou, Dan Xiang, Adrienne H. K. Roeder, Olivier Hamant, Lilan Hong

**Author notes:** These authors contributed equally (Shouling Xu, Xi He).

## Abstract

As in origami, morphogenesis in living systems heavily relies on tissue curving and folding, through the interplay between biochemical and biomechanical cues. In contrast, certain organs maintain their flat posture over several days. Here we identified a pathway, which is required for the maintenance of organ flatness, taking the sepal, the outermost floral organ, in Arabidopsis as a model system. Through genetic, cellular and mechanical approaches, our results demonstrate that global gene expression regulator VERNALIZATION INDEPENDENCE 4 (VIP4) fine-tunes the mechanical properties of sepal cell walls and maintains balanced growth on both sides of the sepals, mainly by orchestrating the distribution pattern of AUXIN RESPONSE FACTOR 3 (ARF3). *vip4* mutation results in softer cell walls and faster cell growth on the adaxial sepal side, which eventually cause sepals to bend outward. Downstream of VIP4, ARF3 works through modulating auxin signaling to down-regulate pectin methylesterase VANGUARD1, resulting in decreased cell wall stiffness. Our work unravels a 3-component module, which relates hormonal patterns to organ curvature, and actively maintains sepal flatness during its growth.

## Introduction

In multicellular organisms, organ morphology exhibits astonishing diversity, especially in three dimensions (Jonsson et al., 2023; Lee et al., 2019; Liang and Mahadevan, 2011; Whitewoods et al., 2020). Understanding the mechanisms controlling three-dimensional morphology of organs is critical for unraveling organ function. The establishment and maintenance of three-dimensional morphology in organs are intricate processes that involve a complex interplay of regulatory mechanisms (Aryal et al., 2020; Huang et al., 2018; Jonsson et al., 2021; LeGoff and Lecuit, 2016). The coordination of cell behaviors such as growth, division and differentiation is crucial to maintain proper development and morphology of organs (Basson, 2012).

Within plants, development takes place in mechanically interconnected tissues, where variations in cell expansion at specific locations result in deformations at the organ level, such as buckling or bending (Coen et al., 2004; Echevin et al., 2019; Fendrych et al., 2016). Differential cellular growth between different parts of the tissue is a pivotal factor in the formation of bends in organs such as apical hooks and roots (Aryal et al., 2020; Huang et al., 2018; Jonsson et al., 2021; LeGoff and Lecuit, 2016). During organ buckling or bending, the plant hormones auxin and the cell wall interplay to affect mechanical properties of tissues, leading to mechanical asymmetry and consequently differential growth (Baral et al., 2021; Ikushima et al., 2008; Jonsson et al., 2021). In recent years, only a few modules have been identified to relate hormones, cell wall effectors, tissue mechanics and organ bending (Baral et al., 2021; Jonsson et al., 2021). This is what we undertook here, using the sepal, the outermost floral organ, as a model system.

We report that mutation of *VERNALIZATION INDEPENDENCE 4* (*VIP4*), a gene encoding a subunit of the global transcription regulator polymerase-associated factor complex (Paf1C), leads to abnormal sepal curvature, through AUXIN RESPONSE FACTOR 3 (ARF3)-dependent downregulation of pectin methylesterase VANGUARD1, affecting cell wall stiffness.

## Results

### Mutations in *VIP4* disrupt three-dimensional sepal morphology

To investigate the genes and molecular mechanisms controlling sepal curvature, we screened (Hong et al., 2016) and ultimately isolated two mutants that had altered sepal curvature (Fig. 1). Wild-type (WT) *Arabidopsis thaliana* (henceforth Arabidopsis) flowers at maturity have quite flat sepals, while flowers of these two mutants tend to have sepals that bend outward (Fig. 1A). Thus, these two mutants were named *abnormal organ morphology 1* (*aom1*) and *abnormal organ morphology 2* (*aom2*). Allelic test showed that *aom1* and *aom2* were alleles, although *aom1* flowers showed a stronger sepal outward bending phenotype than *aom2* flowers. In each mutant, the degree of sepal outward bending varied between flowers. To quantify the sepal phenotypes, we recorded the bending angle of the most out-curved sepal in each flower and used it as the sepal outward bending angle of that flower. The bending angle of an out-curved sepal was defined as the angle between the tangent line on the curved part of the sepal and the central axis of the flower (Fig. 1B). We quantified the sepal outward bending angles of mature flowers. 87.4% of *aom1* flowers had a sepal bending angle over 90°, while only 4.5% of *aom2* flowers had a sepal bending angle over 90° (Fig. 1C).

**Figure 1.**
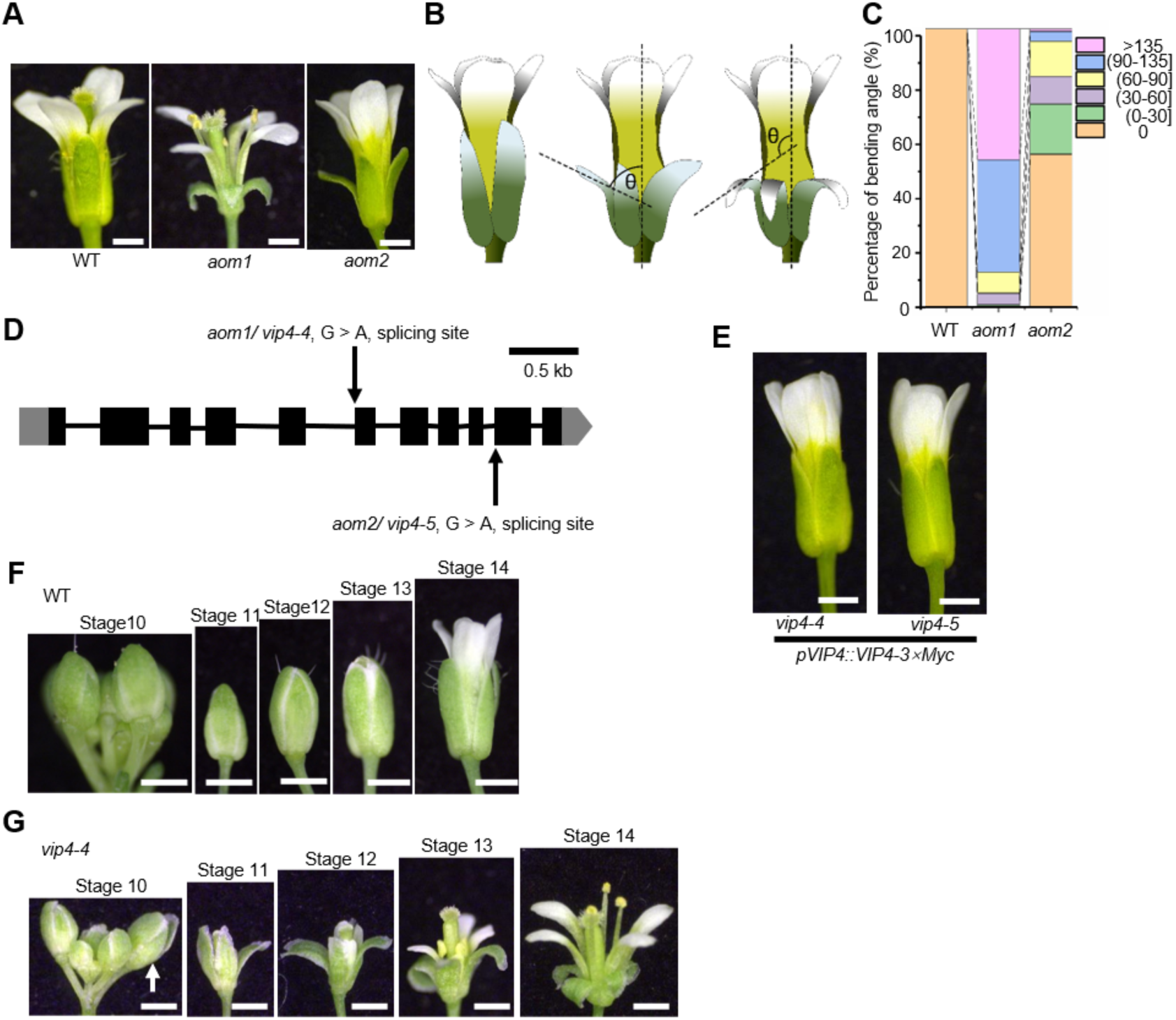
Mutations in the *VIP4* gene lead to sepal outward bending. **(A)** Wild-type (WT), *aom1* and *aom2* flowers at maturity (stage 14). Sepals in *aom1* and *aom2* mutants exhibit outward bending compared to the WT. Scale bars, 0.5 mm. **(B)** Schematic diagram illustrating how to define the bending angle (θ) of a sepal. **(C)** *aom1* displays a stronger sepal bending phenotype compared to *aom2*. The sepal bending angle was quantified for WT and *aom1*, *aom2* based on the criteria shown in panel **(B)**. The bending angle of WT sepal was set to 0. **(D)** Mutation sites of *vip4* alleles isolated in our study. **(E)** T1 plants of *vip4-4* and *vip4-5* transformed with *pVIP4::VIP4-3×Myc*. Scale bars, 0.5 mm. **(F and G**) Developmental progression of sepals in WT and *vip4-4*. The flowers of *vip4-4* **(G)** exhibit normal sepal shape in the early stage of development similar to WT (**F**). *vip4-4* sepals start to bend outward from stage 10. Scale bars, 0.5 mm.

Bulked segregant analysis sequencing (BSA-seq) of *aom1* identified a G-to-A point mutation at a splice acceptor site of *VIP4* (*AT5G61150*), which encodes a crucial component protein of the polymerase–associated factor complex (Paf1C) (Fig. 1D). This point mutation disrupted normal mRNA splicing (Supplemental Fig. S1A) and caused a dramatic decrease in *VIP4* transcription level (Supplemental Fig. S1B). Since *aom1* and *aom2* are allelic, we performed Sanger sequencing on *aom2*, and found that it had a G-to-A point mutation at the splice site between the tenth exon and the ninth intron of *VIP4* (Fig. 1D). Given that *vip4-4* exhibits stronger phenotypes than *vip4-5*, most work in this study was performed on *vip4-4*. Several transfer-DNA (T-DNA) insertion alleles of *VIP4* (*vip4-1*, *vip4-2*, *vip4-3*) have been previously reported(Zhang and van Nocker, 2002). Thus, we renamed *aom1* and *aom2* as *vip4-4* and *vip4-5*, respectively. To verify that mutations in *VIP4* caused sepals to bend outward, the VIP4-3×Myc fusion protein under its endogenous promoter (*pVIP4::VIP4-3×Myc*) was transformed into *vip4-4* and *vip4-5*, and it rescued *vip4-4* and *vip4-5* phenotypes (12/12 and 23/25 rescued in T1, respectively) (Fig. 1E). These results indicate the functionality of the fusion protein and provide additional confirmation of the crucial role of VIP4 in the regulation of sepal curvature.

### *vip4-4* adaxial sepal epidermis has softer cell walls and faster cell growth compared to the abaxial epidermis

We determined when during development the abnormality in mutant sepals first occurred. *vip4-4* flowers had a normal sepal morphology at early stages until stage 10 when their sepals began to bend outward, causing the flowers to fail to close (Fig. 1F and 1G). Thus this phenotype is not the result of defective initiation, but instead emerges from sepal growth later on in development.

In organs such as roots and apical hooks, asymmetric growth on both sides of the organ drives the bending(Aryal et al., 2020; Jonsson et al., 2021; Jonsson et al., 2023). To check whether a comparable growth pattern occurs in sepals, we live-imaged WT and *vip4-4* flowers from stage 10, i.e. when the sepal outward bending started in *vip4-4* (Fig. 2). Both the adaxial and abaxial sides of the same sepal were imaged and the cellular growth rates of both sides were compared. In WT sepals, adaxial cells and abaxial cells grew at similar rates. In contrast, in *vip4-4* sepals, adaxial cells exhibited slightly, but significantly, faster (1.07 ± 0.018 folds, mean ± s.d., n = 3) growth than abaxial cells (Fig. 2A-2C). These results suggest that, over the duration of sepal growth from stage 10 to stage 14 (i.e. 5 days), this small difference would cumulate and explain the outward bending of *vip4* sepals, while WT sepals would maintain symmetric growth and generate unbent sepals.

**Figure 2.**
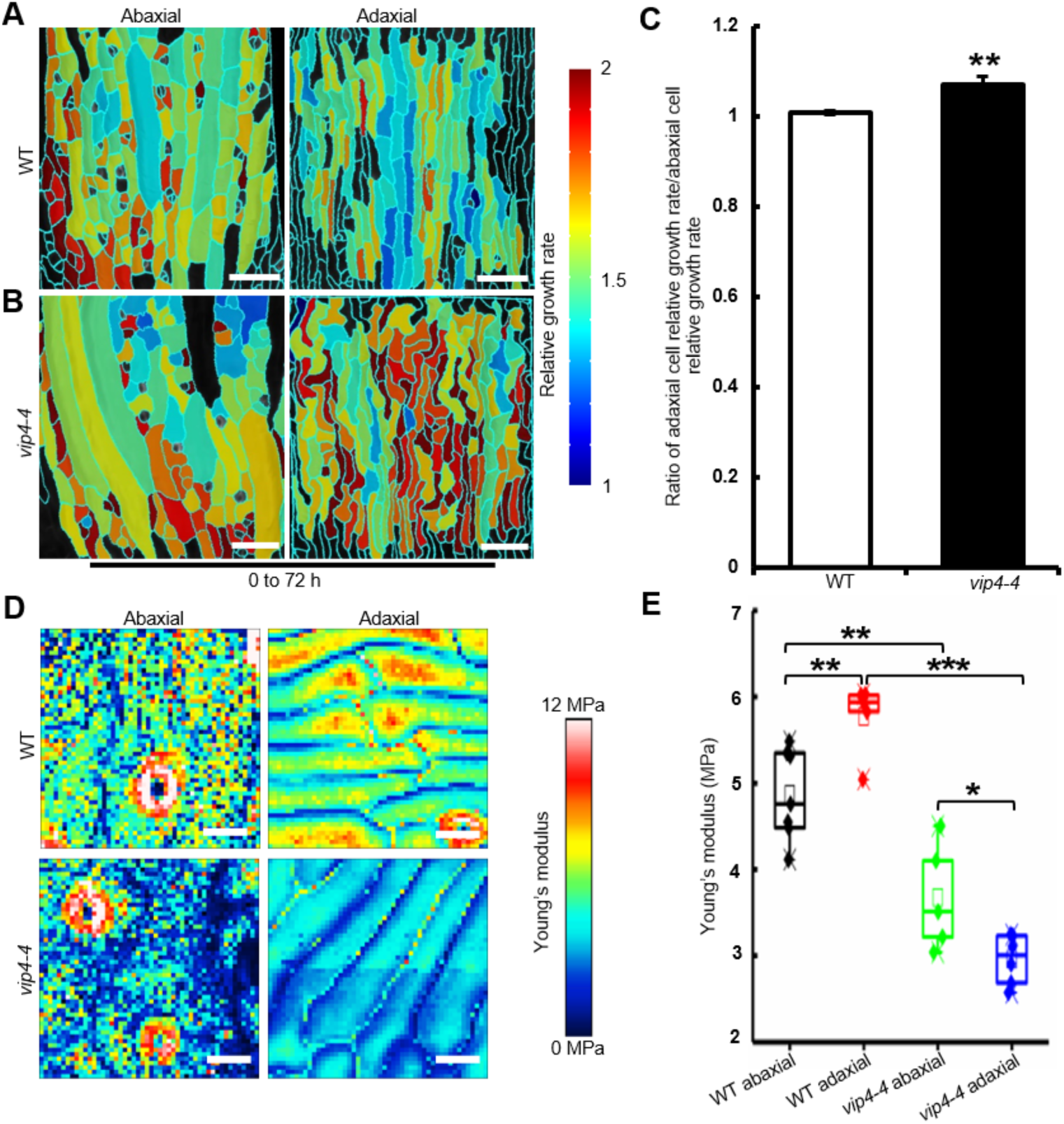
*vip4-4* sepals show differential cell growth and cell wall stiffness. **(A and B)** Heatmaps show 72-hour cellular growth rates on the abaxial (left) side and adaxial (right) side of WT **(A)** and *vip4-4* **(B)** sepals at stage 10. The sepal cellular growth rate was quantified through live imaging. Red and blue colors indicate high and low relative growth rate, respectively. The growth rate is calculated as the ratio of cell area at 72 h to cell area at 0 h. n = 3 biological flowers, all showing similar trends. Scale bars, 50 µm. **(C)** The ratios of adaxial cell relative growth rate to abaxial cell relative growth rate in WT and *vip4-4* sepals. In *vip4-4* sepals, adaxial cells grow faster than abaxial cells, while in WT sepals, adaxial cells and abaxial cells have similar growth rates. Data are mean ± s.d., n = 3 flowers. Two-tailed Student’s t test, \*\**P* < 0.01. Exact *P* value is listed in Supplemental Dataset S6. **(D and E)** *vip4-4* sepals have softer cell wall. **(D)** AFM measurement of cell wall stiffness (Young’s modulus) of both sides for WT and *vip4-4* sepals at stage 10. In the heatmap, stiff points are shown in red and soft points are shown in blue. Scale bars, 10 µm. **(E)** The average apparent elastic modulus calculated from AFM measurements. n = 7 for WT abaxial, n = 5 for WT adaxial, n = 5 for *vip4-4* abaxial, n = 6 for *vip4-4* adaxial. Two-tailed Student’s t test, \**P* < 0.05, \*\**P* < 0.01 and \*\*\**P* < 0.001. Exact *P* values are listed in Supplemental Dataset S6. For the boxplots, the box extends from the lower to upper quartile values of the data, with a line at the median.

During the bending process in Arabidopsis apical hooks, the elongation rates of cells exhibit a correlation with the mechanical properties of their walls (Aryal et al., 2020; Jonsson et al., 2021). To determine whether *vip4-4* exhibits differential cell wall mechanical properties on both sides of its sepals, we used atomic force microscopy (AFM) to quantify cell wall stiffness of the abaxial and adaxial sides of WT and *vip4-4* sepals at stage 10. In WT sepals at stage 10, cell walls on the abaxial side were softer than on the adaxial side (Fig. 2D and 2E). At similar stage, *vip4-4* sepals had significantly softer cell walls on both the abaxial and adaxial sides compared with WT counterparts (Fig. 2D and 2E). Strikingly, *vip4-4* sepals exhibited a significant decrease in cell wall stiffness on the adaxial side compared with the abaxial side, in contrast to the WT. The cellular growth rate and cell wall stiffness in WT and *vip4-4* sepals demonstrate that stiffer walls correlate with slower growth in sepal development, in line with what have been reported in other organs (Aryal et al., 2020; Jonsson et al., 2021; Zhu et al., 2020).

These measurements on cellular growth rate and cell wall stiffness together support the most parsimonious model that in *vip4-4* sepals, softer cell wall contributes to faster cellular growth on the adaxial side, which leads to growth imbalance between the two sides and eventually organ outward bending.

### Ectopic *ARF3* accumulation promotes sepal outward bending in *vip4-4* mutant

Having explored the cellular basis leading to outward bending in *vip4-4* sepals, we next investigated genetic mechanisms involved in VIP4’s controlling of sepal curvature. Paf1C is known to maintain appropriate gene expression by transcription regulation or epigenetic modification of chromatin (Betz et al., 2002; Ng et al., 2003). Mutations in Paf1C components lead to down-regulation/up-regulation of target genes and affect multiple biological processes (Oh et al., 2008). To identify the biological processes that are regulated by VIP4 during sepal morphogenesis, RNA sequencing (RNA-seq) was performed on WT and *vip4-4* sepals at about stage 10. The RNA-seq analysis identified 3714 differentially expressed genes (DEGs) (Supplemental Dataset S1). Gene ontology (GO) term analysis of DEGs revealed an enrichment of biological processes including “cell wall modification”, “response to hormone stimulus” and “pectin metabolic process” (Supplemental Fig. S2A; Supplemental Dataset S2).

One of the DEGs up-regulated in *vip4-4* sepals is *ARF3* (Fig. 3), which has been shown to play a crucial role in the abaxial/adaxial tissue patterning of aerial lateral organs (Andres-Robin et al., 2020; Fahlgren et al., 2006; Garcia et al., 2006; Pekker et al., 2005). We hypothesized that *ARF3* might be involved in formation of the differential growth or differential cell wall stiffness on the abaxial/adaxial sides of *vip4-4* sepals. To test this, we disrupted the up-regulation of *ARF3* in *vip4-4* by crossing *vip4-4* with an *ARF3* mutant *arf3-1* and observed that the mutation of *ARF3* partially suppressed sepal outward bending in *vip4-4* (Fig. 3B and 3C).

**Figure 3.**
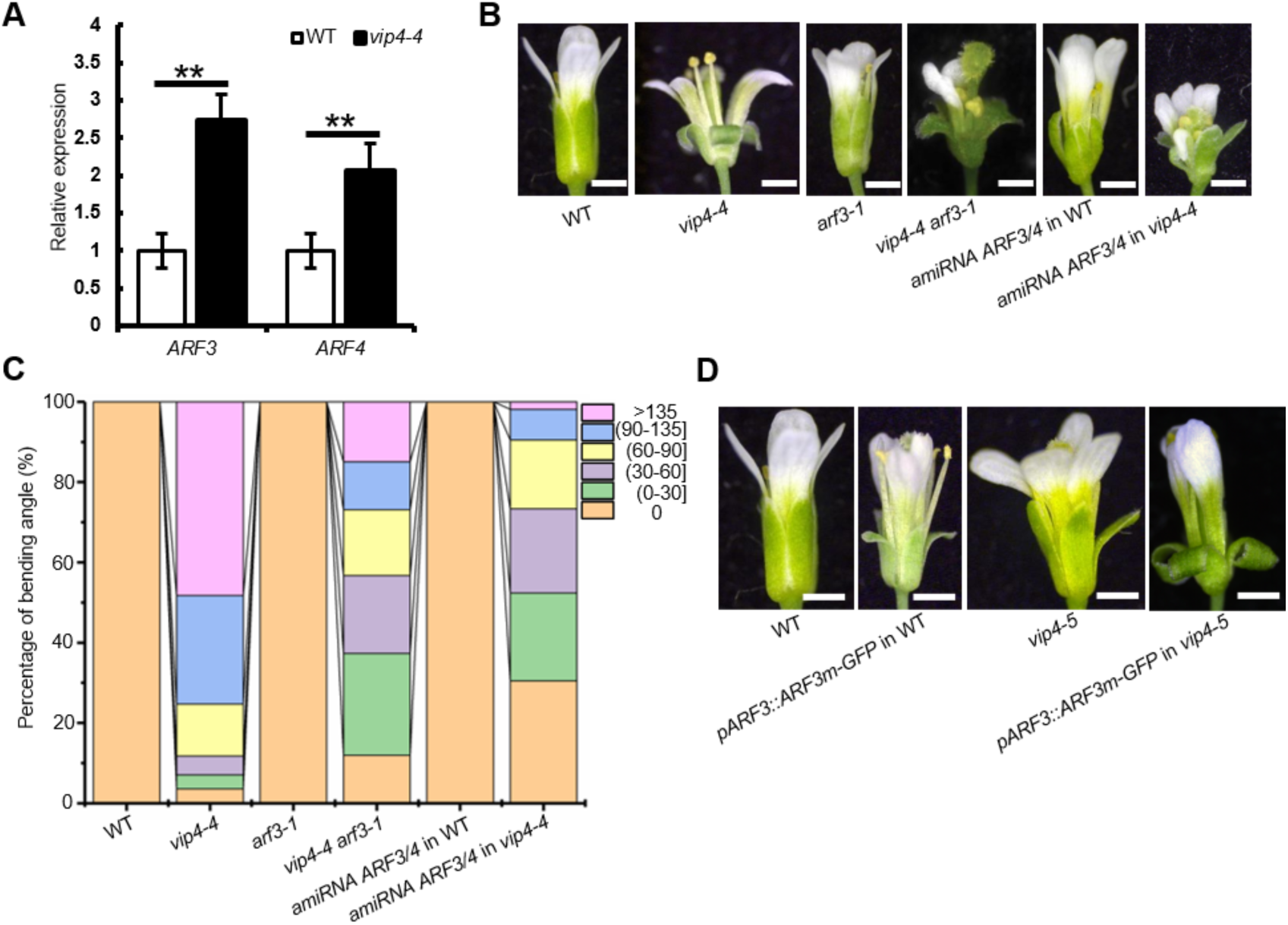
*ARF3* expression levels affect sepal curvature. **(A)** RNA-seq analysis revealing up-regulation of *ARF3* and *ARF4* in *vip4-4* sepals compared to WT. Data are mean ± s.d., n = 3 replicates. Adjusted *P* value (*P*-adj) is obtained via Benjamini-Hochberg correction. \*\**P*-adj < 0.01. Exact *P*-adj are listed in Supplemental Dataset S6. **(B)** Flowers of WT, *vip4-4*, *arf3-1*, *vip4-4 arf3-1*, *amiRNA ARF3/4* in WT and *amiRNA ARF3*/*4* in *vip4-4*. Scale bars: 0.5 mm. **(C)** Quantification of sepal bending angle of the indicated lines, showing reduced sepal bending angles in the *vip4-4* background when *ARF3* was mutated or knocked down. The bending angle of WT sepal was set to 0. **(D)** Flowers of WT, *pARF3::ARF3m-GFP* in WT, *vip4-5*, *pARF3::ARF3m-GFP* in *vip4-5*, showing increased *ARF3* expression level could lead to or enhance sepal outward bending. WT is reproduced from Fig. 2B. Scale bars: 0.5 mm.

The expression of *ARF3* homolog *ARF4* was also up-regulated in *vip4-4* sepals (Fig. 3A and Supplemental Fig. S3A). Considering the similarities in function between *ARF3* and *ARF4*, we speculated that *ARF3* and *ARF4* might be functionally redundant in regulating sepal curvature. Unfortunately, we could not obtain triple mutants of the *VIP4*, *ARF3* and *ARF4* genes, since *VIP4* and *ARF4* are located very close to each other in the genome. To circumvent this problem, we designed *amiRNA ARF3/4*, an artificial RNA that specifically and simultaneously targeted *ARF3* and *ARF4*. Expressing *amiRNA ARF3/4* in WT produced a very similar phenotype to the *ARF3* and *ARF4* double mutant *arf3-1 arf4-2* (Supplemental Fig. S3B), indicating that *amiRNA ARF3/4* worked. Introducing *amiRNA ARF3/4* into *vip4* mutants could significantly suppress sepal outward bending in *vip4-4* and *vip4-5* (Fig. 3B and 3C and Supplemental Fig. S3C). These results suggest that *ARF3* and *ARF4* function redundantly and are necessary in the outward bending of *vip4* sepals.

Since *ARF3* showed higher expression level in WT sepals and more significant up-regulation in *vip4-4* sepals than *ARF4*, we focused on *ARF3* for further study. Having found down-regulation of *ARF3* suppressed the sepal outward bending phenotype in *vip4*, we further investigated whether up-regulation of *ARF3* would promote sepal outward bending. Previous studies showed that *ARF3* expression was regulated by the adaxially expressed mobile *TAS3-*derived trans-acting small interfering RNAs (tasiRNAs) which forms a gradient along the adaxial-abaxial axis and restricts its targets, *ARF3* and *ARF4*, to the abaxial side of the organ (Chitwood et al., 2009; Fahlgren et al., 2006; Garcia et al., 2006; Pekker et al., 2005). We expressed a tasiRNA resistant version of *ARF3* (*pARF3::ARF3m-GFP*) (Liu et al., 2014) to make ARF3 levels less polarized in sepals. Sepal outward bending could be observed in *pARF3::ARF3m-GFP* lines (Fig. 3D), although to a lesser extent compared with *vip4-4*. Since it might not be easy to observe a stronger sepal bending phenotype in *vip4-4*, we introduced *pARF3::ARF3m-GFP* into the weaker allele *vip4-5*, and found the transgene greatly enhanced the *vip4-5* sepal bending phenotype (Fig. 3D). The observations show that ectopic ARF3 accumulation promotes sepal outward bending, and suggest that the asymmetric pattern of ARF3 under the control of VIP4 regulates sepal curvature.

### VIP4 modulates ARF3 distribution

To investigate this scenario, we first asked whether ARF3 is a direct target of VIP4 (Fig. 4). Since Paf1C is recruited to the active Pol II elongation machinery and regulates transcriptional elongation (Betz et al., 2002), we performed a chromatin immunoprecipitation-qPCR (ChIP-qPCR) assay using *pVIP4::VIP4-3×Myc* inflorescences and found that VIP4 directly binds to *ARF3* gene (Fig. 4A).

**Figure 4.**
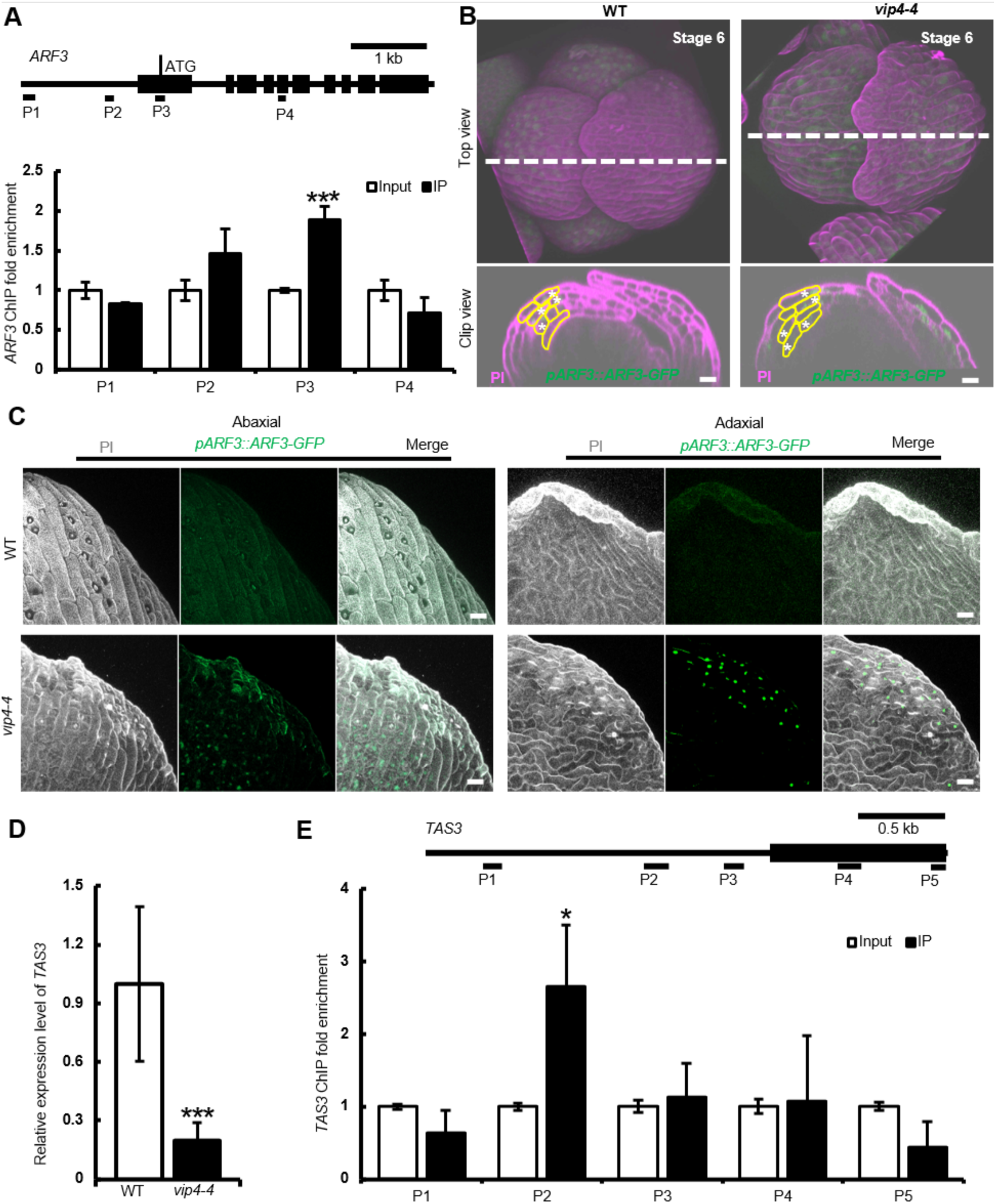
VIP4 regulates *ARF3* expression profile through binding to *TAS3* and *ARF3*. **(A)** Diagrams illustrating the gene structure of *ARF3* with ChIP-qPCR target positions shown and ChIP-qPCR results showing the enrichment of VIP4 in *ARF3* genomic region using anti-Myc antibody. Data are mean ± s.d., n = 3 replicates. Two-tailed Student’s t test, \*\*\**P* < 0.001. Exact *P* value is listed in Supplemental Dataset S6. **(B)** ARF3 distribution pattern in WT and *vip4-4* sepals at stage 6. Confocal images showing the distribution of ARF3 (*pARF3::ARF3-GFP*, green) in WT and *vip4-4* flowers at stage 6 stained with PI (magenta). Upper panels show the front view of the flowers. Lower panels show the side view of clips at the white lines in upper panels. *ARF3* is expressed in the abaxial three layers of cells in WT sepals, while in *vip4-4* sepals ARF3 is distributed in all the four layers of cells. Some cells in the transverse sections are outlined in yellow to show the four layers of sepal cells. Asterisks mark ARF3-GFP signals. Scale bars, 20 µm. **(C)** ARF3 distribution pattern in WT and *vip4-4* sepals at stage 10. ARF3-GFP signals could not be detected on either sides of sepals at stage10 in WT, while remain visible in *vip4-4* sepals at the same stage (some ARF3-GFP signals highlighted with red circles). Gray, propidium iodide (PI) staining the cell wall; green, *pARF3::ARF3*-GFP. Scale bars, 20 µm. n= 10 flowers for WT. n=8 flowers for *vip4-4*. **(D)** RNA-seq data showing downregulation of *TAS3* in *vip4-4* sepals compared to WT. Data are mean ± s.d., n = 3 replicates. *P*-adj is obtained via Benjamini-Hochberg correction. \*\*\**P*-adj < 0.001. Exact *P*-adj is listed in Supplemental Dataset S6. **(E)** Diagrams illustrating the gene structure of *TAS3* with ChIP-qPCR target positions shown and ChIP-qPCR results showing the enrichment of VIP4 in *TAS3* promoter using anti-Myc antibody. Data are mean ± s.d., n = 3 replicates. Two-tailed Student’s t test, \**P* < 0.05. Exact *P* value is listed in Supplemental Dataset S6.

Second, we checked the localization of ARF3 in WT and *vip4-4* sepals using the *pARF3::ARF3-GFP* (Liu et al., 2014) reporter. We found that at early developmental stages, ARF3 was primarily located to the abaxial side of WT sepals, resembling its distribution pattern in leaves (Burian et al., 2022). In *vip4-4* sepals at similar stages, the distribution of ARF3 extended towards the adaxial side (Fig. 4B). The ectopic expression of *ARF3* in *vip4-4* sepals lasted to stage 10, when ARF3 was detected on both sides of *vip4-4* sepals, whereas *ARF3* expression could not be detected on either side of WT sepals at this stage (Fig. 4C). In short, *ARF3* expression in *vip4-4* sepals shows prolonged duration and an expanded domain. We also noticed that WT plants transformed with *pARF3::ARF3-GFP* had normal sepal morphology, while ectopic expression of *ARF3* in WT with *pARF3::ARF3m-GFP* generates sepal outward bending, which is consistent with our scenario in which sepal outward bending in *vip4-4* is mainly caused by ectopic ARF3 distribution.

As mentioned above, *TAS3* tasiRNAs target *ARF3* and *ARF4*. In our study, we observed a significant decrease in the expression of *TAS3* in *vip4-4* sepals compared to WT (Fig. 4D). The down-regulation of *TAS3* may attenuate *TAS3*’s repression of *ARF3* expression and lead to expanded activity of *ARF3* from the abaxial to the adaxial side of *vip4-4* sepal. ChIP-qPCR showed VIP4 could bind the *TAS3* promoter regions (Fig. 4E). Taken together, these results suggested that VIP4 could regulate ARF3 distribution profile directly by binding to *ARF3* and indirectly by affecting *TAS3* expression.

### ARF3 acts through auxin signaling in regulating sepal morphology

In the simplest scenario, ARF3 would regulate sepal curvature through auxin signaling. To test this hypothesis, we checked whether auxin signaling is required for the outward bending of *vip4-4* sepals (Fig. 5). Auxin signaling is inhibited in the auxin receptor quadruple mutant *tir1-1 afb1-1 afb2-1 afb3-1* (Dharmasiri et al., 2005). We generated a *vip4-6 tir1-1 afb1-1 afb2-1 afb3-1* pentuple mutant by knocking out *VIP4* via CRISPR-Cas9 (Supplemental Fig. S4A) in *tir1-1 afb1-1 afb2-1 afb3-1* mutant and found that the sepals of the pentuple mutant did not bend outward (Fig. 5A), suggesting that functional auxin signaling is necessary in the outward bending process of *vip4* sepals.

**Figure 5.**
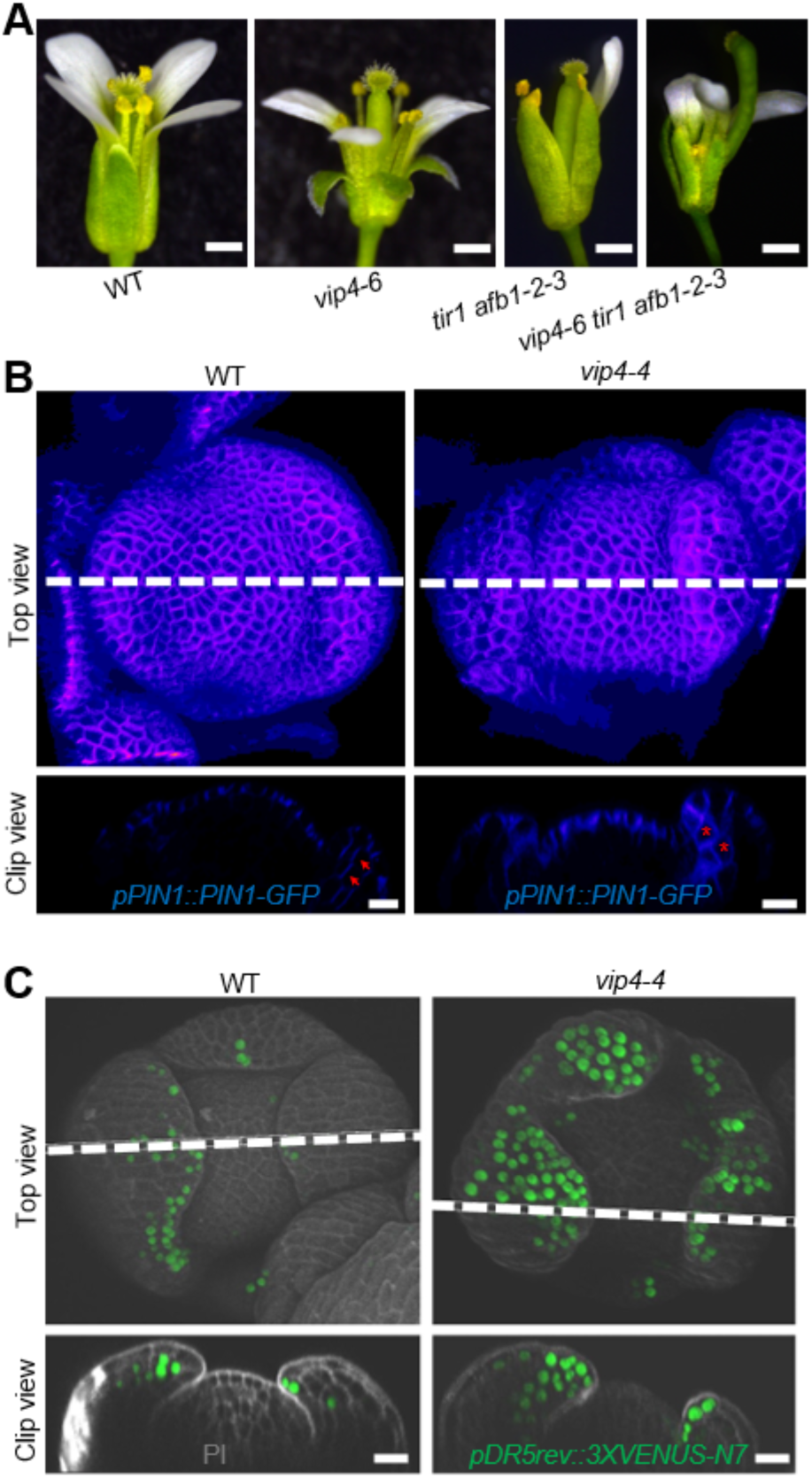
Auxin signaling is involved in the outward bending of *vip4-4* sepals. **(A)** Flowers of WT, *vip4-6*, *tir1 afb1-2-3*, and *vip4-6 tir1 afb1-2-3*. Inhibiting auxin signaling in *vip4* suppresses sepal outward bending. Scale bars: 0.5 mm. **(B)** Confocal images showing PIN1 (*pPIN1::PIN1-GFP*, blue) distribution in WT and *vip4-4* flowers at stage 4. Upper panels show the front view of the flowers. Lower panels show the side view of clips at the white lines in upper panels. Cells in WT exhibit PIN1-GFP polarity from the base to the tip of the sepal (as the red arrows point), while cells in *vip4-4* sepals show no polarized localization of PIN1-GFP (marked with asterisks). Scale bars, 20 µm, n= 5 flowers. LUT is gem in ImageJ. **(C)** Confocal images showing auxin distribution (*pDR5rev::3×VENUS-N7*, green) in WT and *vip4-4* flowers at stage 5 stained with PI (gray). Upper panels show the front view of the flowers. Lower panels show the side view of clips at the white lines in upper panels. DR5 signals accumulate at the tips of four sepals in WT, while have a broader distribution in *vip4-4* sepals. Scale bars, 20 µm. n = 5 flowers.

Previous studies revealed that ARF3 influences auxin distribution through regulating the expression of auxin transporter genes (Simonini et al., 2017). There were also significant changes observed in the expression of auxin transporter genes in *vip4-4* (Supplemental Fig. S4B). The distribution of auxin in the shoot is generally determined by the polar localization of the auxin efflux carrier PIN-FORMED1 (PIN1) (Heisler et al., 2005). ARF3 has been reported to bind to the *PIN1* genomic region directly and promotes the expression of *PIN1* in previous research (Simonini et al., 2017). *PIN1* was up-regulated in *vip4-4* sepals compared with WT (Supplemental Fig. S4B). It is likely that the altered distribution of ARF3 in *vip4-4* sepals affects PIN1 localization. We checked PIN1 localization pattern in WT and *vip4-4* sepals using *pPIN1::PIN1-GFP* reporter. Cells in WT exhibited PIN1-GFP polarity from the base to the tip of the sepal, whereas cells in *vip4-4* sepals showed no polarized localization of PIN1-GFP (Fig. 5B).

To visualize whether the altered PIN1 localization in *vip4-4* sepals disrupts normal auxin distribution, we examined auxin activity pattern in WT and *vip4-4* sepals using the auxin response reporter *pDR5rev::3×VENUS-N7* (*DR5*) (Heisler et al., 2005; Zhu et al., 2020) . *vip4-4* sepals had broader auxin activity compared to WT at early developmental stages (Fig. 5C and Supplemental Fig. S4C and 4D). In WT, DR5 signal accumulated mainly in the inner tissue of sepals at stage 5. In contrast, in *vip4-4* sepals at similar stages, DR5 signal displayed an expanded distribution domain, with strong signals visible in all cell layers (Fig. 5C). Although neither WT nor *vip4-4* sepals had detectable *DR5* activities at late developmental stages, the expression levels of numerous genes associated with auxin synthesis, signaling, and response were up-regulated in *vip4-4* sepals at stage 10 compared with WT (Supplemental Fig. S4E). Collectively, these data indicate that *vip4-4* sepals have broader and increased auxin levels.

A plethora of studies have demonstrated that auxin influences the mechanical properties of cell walls, although the exact influence varies among tissues (Fendrych et al., 2016; Jonsson et al., 2021; Velasquez et al., 2021; Zhu et al., 2020). To investigate the impact of auxin on the mechanical properties of sepal cell walls, we treated WT flowers at stage 10 with indole-3-acetic acid (IAA) and performed osmotic treatment to assess cell wall stiffness in sepals. We found that WT sepals treated with IAA had a greater cell wall shrinkage compared to the mock treated sepals (Supplemental Fig. S5A and 5B), indicating that increased auxin levels softens sepal cell walls. This finding fits with the increased auxin levels and the decreased cell wall stiffness in *vip4-4* sepals. Our results are also consistent with previous research in sepal primordia, which reported that sepal primordia with weaker auxin signaling had stiffer cell walls (Zhu et al., 2020).

Altogether, our data indicate that in *vip4-4* sepals, ectopic ARF3 distribution leads to increased and ectopic auxin accumulation which consequently reduce the stiffness of sepal cell walls, disrupting the balanced growth between adaxial and abaxial sides and affecting sepal curvature.

### ARF3 regulates sepal curvature through regulating the expression of pectin modification enzymes

Previous studies have demonstrated that ARF3 affects the expression of genes encoding pectin methylesterases (Andres-Robin et al., 2018). In our RNA-seq data, we found many genes differentially expressed in *vip4-4* sepals were related to pectin modification (Supplemental Fig. S2B), which prompted us to hypothesize that the ectopic expression of *ARF3* promotes sepal outward bending not only through modulating auxin distribution, but also through regulating the expression of pectin modification enzymes (Fig. 6). Among the *vip4-4* DEGs encoding pectin methylesterases, the *VANGUARD1* (*VGD1*, *AT2G47040*) gene exhibited a very high expression level in WT sepals and a significant down-regulation in *vip4-4* sepals (Fig. 6A and Supplemental Fig. S2B; Supplemental Dataset S3). Studies in pollen tubes showed that *VGD1* potentially contributed to cell wall stiffening (Jiang et al., 2005). In seedlings, *VGD1* was down-regulated shortly after auxin treatment (Supplemental Fig. S6A). In view of these, we investigated whether VGD1 works downstream of ARF3 and auxin in modulating sepal curvature.

**Figure 6.**
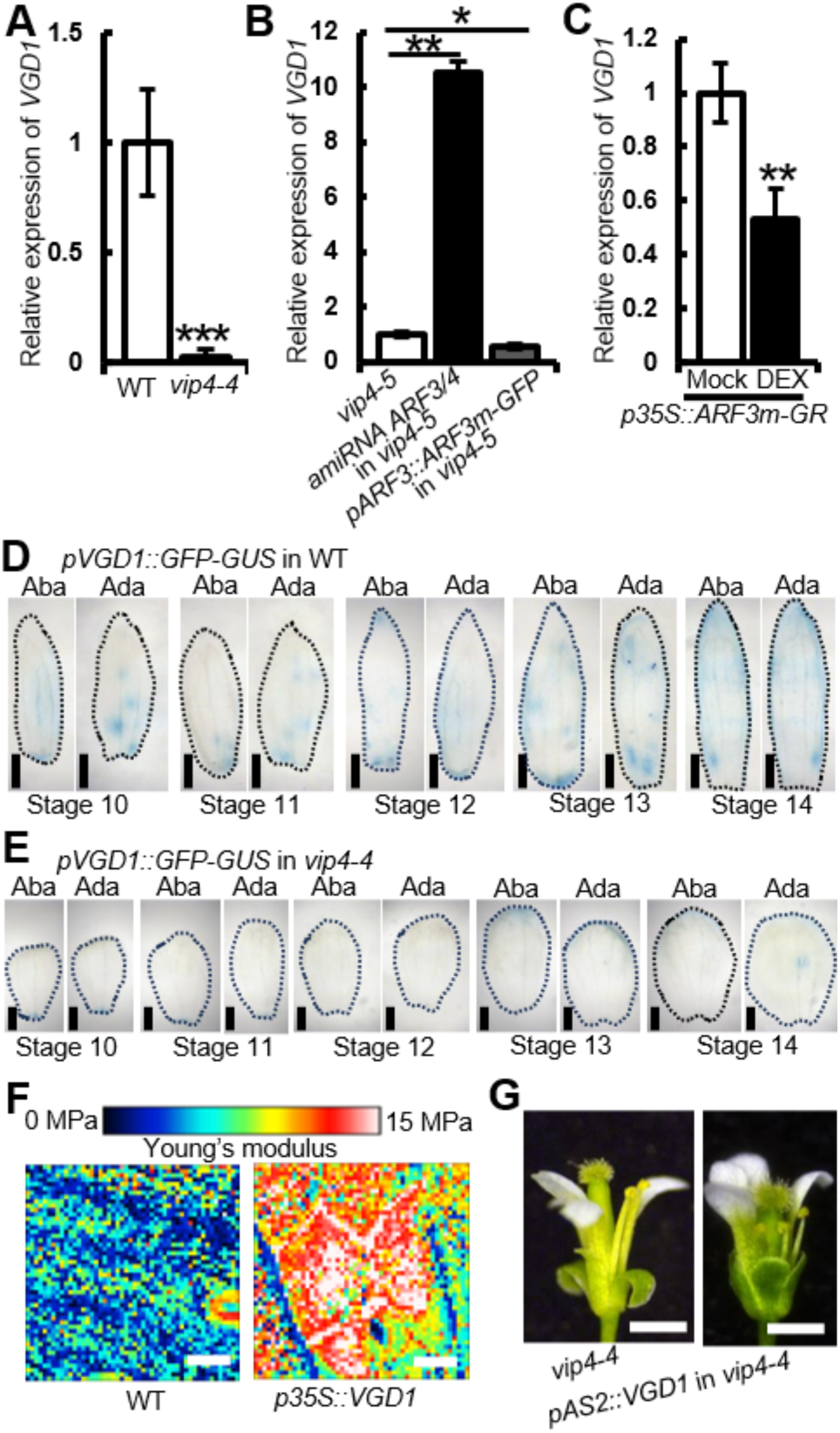
ARF3 may affect cell wall stiffness by regulating *VGD1* expression. **(A)** RNA-seq analysis revealing down-regulation of *VGD1* in *vip4-4* sepals compared to WT. Data are mean ± s.d., n = 3 biological replicates. *P*-adj is obtained via Benjamini-Hochberg correction. *** *P*-adj < 0.001. Exact *P*-adj is listed in Supplemental Dataset S6. **(B and C)** *ARF3* negatively regulates *VGD1* expression. **(B)** *VGD1* expression is up-regulated by *amiRNA ARF3*/*4* while down-regulated by *ARF3* overexpression (*pARF3::ARF3m-GFP*) in *vip4-5* background. **(C)** DEX-treated *p35S:ARF3m-GR* flowers showing down-regulation of *VGD1* in contrast to solvent DMSO-treated flowers. Data are mean ± s.d., n = 3 biological replicates. Two-tailed Student’s t test, \**P* < 0.05, \*\**P* < 0.01. Exact *P* values are listed in Supplemental Dataset S6. **(D and E)** *VGD1* expression profiles during late development stage in WT **(D)** and *vip4-4* **(E)** sepals. From stage 10 to stage 14, WT shows higher *VGD1* expression level on both sides of the sepals compared to *vip4-4* (sepals outlined with dashed lines). Scale bars, 200 µm. **(F)** Overexpression of *VGD1* stiffens cell walls in sepals. AFM measurements show increased cell wall stiffness in *p35S:VGD1* lines. Scale bars, 10 µm. **(G)** *pAS2::VGD1* suppresses the *vip4-4* sepal outward bending by hardening the adaxial cell wall. Scale bars, 0.5 mm.

First, we checked how the expression of *VGD1* was affected by *ARF3* levels. Since *vip4-5* plants with varied *ARF3/4* levels exhibited a positive correlation between *ARF3*/*4* levels and the extent of sepal bending, we analyzed *VGD1* expression in these *vip4-5* plants. Flowers of *amiRNA ARF3/4 vip4-5* had a significant up-regulation of *VGD1*, while *pARF3::ARF3(m)-GFP vip4-5* flowers had a decreased level of *VGD1* expression, compared with *vip4-5* flowers (Fig. 6B). The expression of *VGD1* was also up-regulated in *arf3-1* inflorescences compared to WT ones (Supplemental Fig. S6B). Together these data reveal a negative correlation between the expression levels of *ARF3* and *VGD1*. To further explore how ARF3 regulated *VGD1*, we generated transgenic plants containing a dexamethasone (DEX) inducible construct *p35S:ARF3m-GR* that expressed the tasiRNA resistant version of ARF3-GR (ARF3m-GR) fusion protein. In *p35S:ARF3m-GR*, the *ARF3m* cDNA was fused translationally to a DNA fragment encoding the hormone binding domain of the glucocorticoid receptor and placed under the constitutive 35S promoter of the cauliflower mosaic virus. After the DEX treatment, *VGD1* expression in *p35S:ARF3m-GR* flowers was down-regulated (Fig. 6C), indicating that ARF3 represses *VGD1* expression.

To study how the up-regulated and ectopic expression of *ARF3* in *vip4-4* sepals affect *VGD1* expression in detail, we examined the tissue expression pattern of *VGD1* gene. We introduced a *pVGD1::GFP-GUS* reporter into WT and *vip4-4* plants. However, no visible GFP signal was detected in either WT or *vip4-4* sepals with the *pVGD1::GFP-GUS* transgene. Therefore, we focused on GUS signals. We compared GUS staining in WT and *vip4-4* flowers ranging from stage 10 to stage 14 during which stages the sepal outward bending phenotype started and intensified in *vip4-4*. From stage 10 to stage 14, WT had higher *VGD1* expression on both sides of the sepals than *vip4-4* (Fig. 6D and 6E), which together with the fact that *ARF3* was up-regulated and ectopically expressed in *vip4-4* sepals further support the preceding conclusion that *VGD1* expression is negatively regulated by *ARF3*.

*VGD1* has been suggested to stiffen the cell wall in pollen tubes (Jiang et al., 2005). To check how the alteration of *VGD1* expression might affect sepal cell wall mechanical properties as well as sepal morphology, we increased the *VGD1* levels in WT by generating a *VGD1* overexpression line *p35S::VGD1*. The *p35S::VGD1* plants had darker leaves, slower vegetative growth, and smaller sepals compared to WT (Supplemental Fig. S7A and 7B). Cell wall stiffness measurement using AFM showed that *p35S::VGD1* sepals had higher cell wall stiffness compared to WT sepals, indicating VGD1 could lead to stiffened cell walls in sepals (Fig. 6F and Supplemental Fig. S6C). Consequently, VGD1 was used to differentially modify cell wall stiffness in sepals. *VGD1* expressed under an adaxial specific *ASYMMETRIC LEAVES2* (*AS2*) (Xu et al., 2008) promoter (*pAS2::VGD1*) in *vip4-4* well suppressed the outward bending of *vip4-4* sepals (Fig. 6G), confirming that the softer cell wall on the adaxial epidermis contributes to sepal outward bending in *vip4-4*.

Overall, these results provide a complete scenario in which VIP4 directly and indirectly represses ARF3 activity and pattern in sepals, resulting in modified pectin methylesterification and stiffer walls. This ultimately results in a balanced growth rate between adaxial and abaxial sides to maintain flat sepals. Modifications in any of these actors lead to sepal outward bending.

## Discussion

As previously shown in leaves, maintaining organ flatness does not occur by default. It notably requires controlled growth at the organ margins (Nath et al., 2003). Here we exerted this finding to the entire organ, in the case of the sepal: maintaining unbent sepals for several days requires molecular and mechanical control of cell growth rate.

In this work, through studying Arabidopsis mutants with abnormal sepal outward bending phenotype, we discovered that a generic transcription regulator VIP4 is important for maintaining proper sepal morphology. VIP4 regulates sepal morphogenesis mainly through orchestrating the distribution pattern of ARF3 in sepals, while ARF3 in turn modulates auxin signaling and cell wall modification. In WT sepals, with proper VIP4 and ARF3 function, the mechanical properties of sepal cell walls are fine-tuned to achieve balanced tissue growth, and the sepal three-dimensional morphology is maintained. In *vip4* sepals, *ARF3* is ectopically expressed, resulting in altered auxin distribution and cell wall modification, which leads to differential cell wall mechanical properties and consequently differential growth on two sides of the sepals, hence generating outward bent sepals (Fig. 7). Our work reveals that auxin through cell wall mechanics regulate sepal morphogenesis in Arabidopsis and unravels a 3-component module (VIP4-ARF3-VGD1) that coordinates auxin and cell wall mechanics in sepal development.

**Figure 7.**
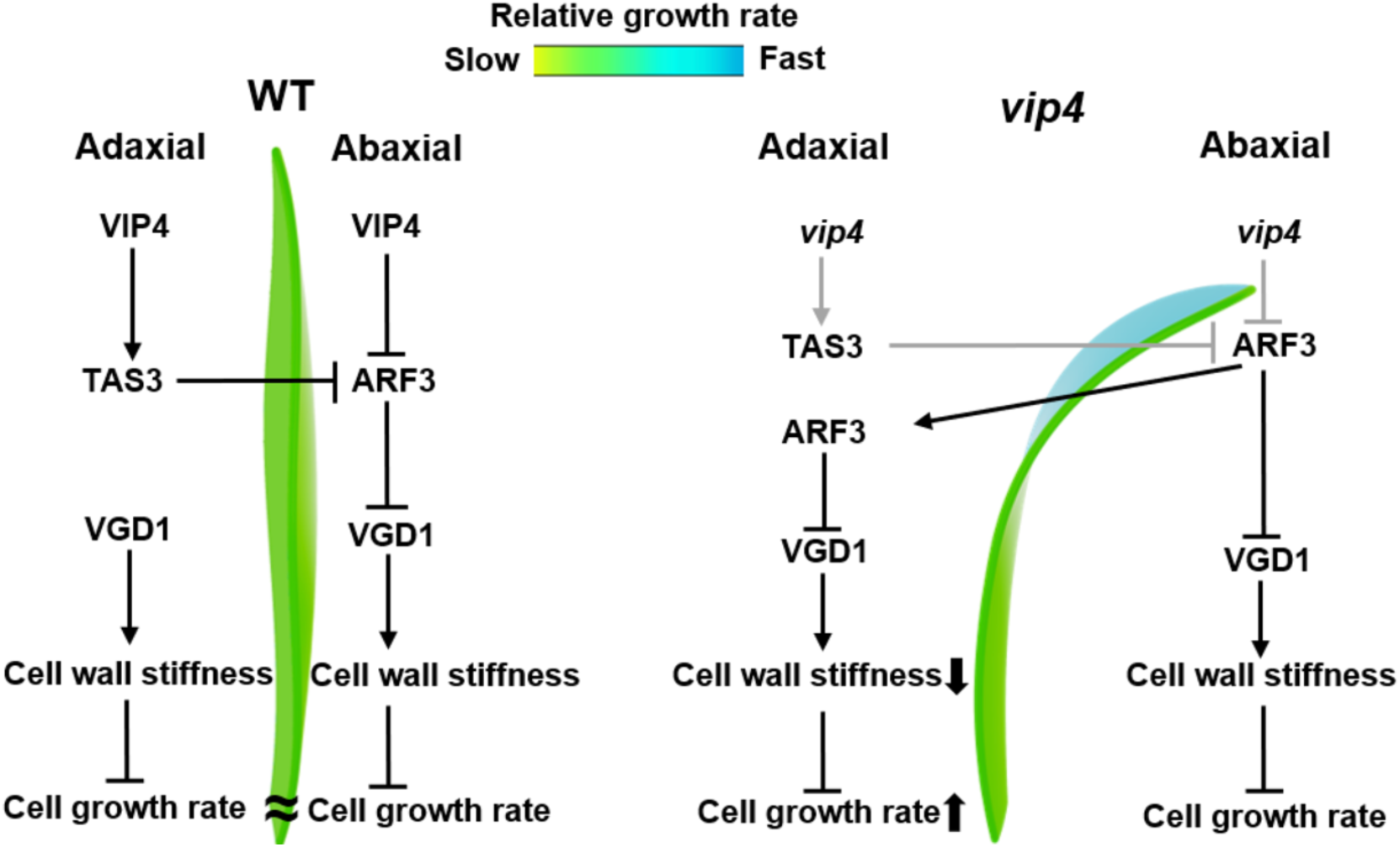
Summary model: a 3-component module controls sepal curvature. In WT sepals, ARF3 expression is restricted to the adaxial side of the sepals due to the presence of functional VIP4 and appropriate expression level of *TAS3*. The proper distribution of ARF3 plays a crucial role in regulating cell wall stiffness by influencing the expression level of *VGD1*. This regulation, in turn, contributes to maintaining balanced tissue growth on both sides of the WT sepal, resulting in a flat sepal structure. However, in *vip4-4* sepals with a mutated *VIP4* gene, there is an up-regulation of *ARF3* expression and a down-regulation of *TAS3* expression. This leads to the ectopic expression of *ARF3*, disrupting its normal adaxial confinement. The abnormal distribution of ARF3 results in an accelerated cell growth rate with softer cell walls by mediating *VGD1* expression. Consequently, this abnormal growth pattern leads to the outward bending of the sepal.

Similar phenotypes have been reported in other Paf1C mutants (He et al., 2004; Trinh et al., 2023; Zhang and van Nocker, 2002), whereas the precise molecular and cellular mechanisms responsible for their bending remained unidentified. Paf1C mutants also exhibit morphological defects in other organs such as cotyledons, siliques, leaves and shoot apical meristems (Fal et al., 2019; Fal et al., 2017; Trinh et al., 2023), indicating that Paf1C plays important roles in regulating the three-dimensional morphology of organs. Paf1C is involved in transcriptional regulation. Mutations in the Paf1C genes lead to dysregulation of global gene expression and subsequent numerous phenotypic abnormalities (Fal et al., 2019; Fal et al., 2017; He et al., 2004; Oh et al., 2008; Trinh et al., 2023; Zhang and van Nocker, 2002; Zhao et al., 2005). Accordingly, we have identified thousands of genes differentially expressed in *vip4* sepals.

Among these genes, *ARF3* exerts a primary role in the outward bending of sepals in *vip4*. *ARF3* expression is regulated at the transcriptional, post-transcriptional and epigenetic levels (Chandler, 2016). In sepals, VIP4 regulates *ARF3* expression at least on two levels: directly by binding to *ARF3* genomic region and indirectly by affecting *TAS3* expression. Via this multifaceted regulation, VIP4 ensures a relatively stable ARF3 distribution pattern, and consequently robust sepal morphogenesis. Paf1C has been implicated in controlling transcriptional noise in Arabidopsis (Ansel et al., 2008; Fal et al., 2019; Trinh et al., 2023). Our data suggests a possible mechanism of Paf1C maintaining the robustness of downstream gene expression.

*ARF3* expression can be induced by exogenous auxin (Zhang et al., 2022) and ARF3 modulates auxin distribution via regulating the expression of polar auxin transportation genes (Simonini et al., 2017). In addition, ARF3 acts as an auxin signaling component (Kuhn et al., 2020; Simonini et al., 2016; Simonini et al., 2018). In view of VIP4’s direct regulation of ARF3 distribution and the altered distribution patterns of auxin and auxin polar transporters in *vip4* sepals, we speculate that ARF3 affects sepal morphogenesis through orchestrating auxin distribution, in line with *ARF3*’s role in auxin signaling and pattern specification (Simonini et al., 2017).

We cannot rule out the possibility that VIP4 affects auxin distribution through directly regulating the expression of genes pertinent to auxin dynamics, or that auxin signaling exerts impacts on sepal morphology through other ARF3/4 independent pathways. We also cannot rule out the possibility that VGD1, with such a striking effect when overexpressed, triggers compensatory effects on cell wall properties, given the size of the PME family and the complexity of the cell wall (Francoz et al., 2019; Hocq et al., 2017).

Although our study focused on the relationship between *ARF3* and genes related to pectin methylesterification, *ARF3* also affects the expression level of genes involved in other cell wall modifications (Supplemental Dataset S3). Numerous studies have demonstrated that auxin activity and cell wall mechanics are highly interrelated in organ morphogenesis (Aryal et al., 2020; Baral et al., 2021; Jonsson et al., 2021; Qi et al., 2017). Our work revealed that auxin and cell wall mechanics also collaborate in shaping three-dimensional sepal morphology, with *ARF3* and its upstream regulator Paf1C acting as overarching coordinators.

## Methods

### Mutations

In this study, Arabidopsis accession *Col*-0 plants were used as wild type (WT). WT seeds were mutagenized with 0.3% (v/v) ethyl methanesulfonate in 10 mL 0.02% (v/v) Tween 20 for 24 hours. M2 plants were examined under a dissecting microscope for the abnormal three-dimensional sepal phenotype. The *vip4-4* and *vip4-5* mutants were back-crossed to *Col*-0 three times to segregate unrelated mutations before further characterization. The *vip4-4* mutated gene was identified through BSA-seq following the standard procedure described in ref. (Takagi et al., 2013). The *vip4-4* mutant has a G-to-A mutation at the junction between the fifth intron and sixth exon within the *VIP4* (*AT5G61150*) gene, which disrupts normal mRNA splicing. The *vip4-5* mutant has a G-to-A mutation at the splice junction between the ninth intron and tenth exon. Allelism tests were conducted between *vip4-4* and *vip4-5*. The F1 displayed the *vip4-4* mutant phenotypes, indicating that these two mutants are allelic.

### Flower staging

Flowers were staged according to ref. (Smyth et al., 1990).

### Live imaging of sepal growth

Live imaging of both sides of sepals and shoot apical meristems was conducted following the sample preparation method outlined in ref. (Stanislas et al., 2017). Briefly, young inflorescences containing different genetic constructs, namely *pUBQ10::Lti6b-tdTomato*, *pDR5rev::3×VENUS-N7* (Heisler et al., 2005; Zhu et al., 2020), *pPIN1::PIN1-GFP*, and *pARF3::ARF3-GFP* (Liu et al., 2014) were dissected at the desired developmental stages using tweezers. For imaging the adaxial and abaxial sides of sepals, all flower buds beyond stage 10 were dissected off and one sepal from flower buds at stage 10 was selectively kept for imaging, while smaller flower buds were retained for further observations. The dissected inflorescences were then cultured in the apex culture medium (containing 1% sucrose, 0.25×vitamin mix and 1% agar) as described in ref. (Stanislas et al., 2017). For the analysis of sepal growth, both the adaxial and abaxial sides of the same sepal were imaged on the first and fourth days, respectively. In cases where the size of the sepals exceeded the image acquisition frame, multiple tiles were stitched together using the Stitching plugin in ImageJ (Preibisch et al., 2009).

All confocal images were acquired using an SP8 laser-scanning confocal microscope (Leica) equipped with a long-distance 25× (NA 0.95) water-dipping objective and a resonant scanner module with a 1.0 µm z-step, except for the imaging of *pARF3::ARF3-GFP*, which was performed using an upright 980 laser-scanning confocal microscope (Zeiss). Excitation and emission wavelengths for fluorescent proteins are indicated in Supplemental Dataset S4.

### Transgenic plants

Otherwise stated, the amplification of PCR fragments was performed using KOD DNA polymerase (Toyobo, CAT KOD-101) and all vectors were generated using T5 exonuclease-dependent assembly (Xia et al., 2019).

To generate *pVIP4::VIP4-3×Myc*, the *VIP4* sequence from the promoter (2036 bp upstream the start codon) to the sequence before the stop codon was amplified from *Col*-0 genomic DNA. The PCR product was cloned into *pRGEcMyc* (He et al., 2018) digested with the restriction enzymes *Pst*I and *Spe*I.

The 4.1-kb upstream promoter and part of the first exon region of the *AS2* gene were first PCR amplified from WT genomic DNA and cloned into *pENTR/D-TOPO* vectors (Invitrogen) as described in the manual. The resultant vector was LR recombined into the gateway vector *pBGFWG2* (Karimi et al., 2002) to generate the final construct *pAS2::GFP-GUS*.

To generate *pVGD1::GFP-GUS*, the *VGD1* promoter (2137 bp upstream of the start codon) was amplified and was inserted into *pAS2::GFP-GUS* digested with *Xba*I and *Pme*I.

To generate *p35S::VGD1* and *pAS2::VGD1*, the *VGD1* sequence from the start codon to the stop codon was amplified from WT genomic DNA. The PCR product was cloned into *pCAMBIA1300-35S* (a binary vector derived from *pCAMBIA1300,* containing the 2× CaMV *35S* promoter and the CaMV terminator) digested with *Kpn*I and *Pst*I and *pAS2::GFP-GUS* digested with *Nco*I and *Asc*I, respectively.

*ARF3*/*ARF4* knockdown mutants were created through the utilization of an artificial microRNA (*amiRNA*) based knockdown approach (Schwab et al., 2006). To design the *amiRNA* constructs, the Web MicroRNA Designer program available at http://wmd3.weigelworld.org/ was employed to concurrently target both *ARF3* and *ARF4*. The *amiRNA* construct was constructed with the endogenous *miRNA319a* as the backbone. The final products were cloned into *pCAMBIA1300-35S* digested with *Kpn*I and *Pst*I.

For the overexpression of *ARF3m*-*GR*, *ARF3m*-*GR* was amplified using *pARF3::ARF3m-GR* (Liu et al., 2014) as a template and then cloned into *pCAMBIA1300-35S* digested with *Kpn*I and *Pst*I.

To generate *VIP4* mutation via CRISPR-Cas9, two guide sequences targeting *VIP4* were designed and then inserted into *pHEE401* vector, according to the guide of ref. (Xing et al., 2014).

All primers used to construct the vectors are listed in Supplemental Dataset S5. All of the final constructs including *pARF3::ARF3-GFP* and *pARF3::ARF3m-GFP* (Liu et al., 2014) were verified by sequencing and transformed into the corresponding plants by *Agrobacterium*-mediated floral dipping. Note that, *pVGD1::GFP-GUS* and *pARF3::ARF3-GFP* were transformed into F1 plants (the hybrid of *vip4-4* and WT) . All T1 plants were grown in plates with corresponding antibiotic resistance (Kanamycin, Hygromycin and Basta). The surviving plants or plants with elongated hypocotyls were then checked for sepal phenotypes or fluorescence at the microscope.

### RNA-seq

For sample collection, the WT and *vip4-4* sepals at stage 10 were dissected under a stereomicroscope and collected into Eppendorf tubes with liquid nitrogen. Three biological replicates were collected, each replicate combining sepals from different individual plants. RNA extraction was done using a commercial kit (Easy Plant RNA Kit, Easy-Do, CAT DR0406050) following the manual. Library preparation was carried out, and their quality was assessed with an Agilent 2100 Bioanalyzer (Agilent Technologies). Subsequently, sequencing was performed on a HiSeq 2500 (Illumina) in accordance with the manufacturer’s guidelines. Raw reads were subjected to cleaning and alignment to the TAIR10 Arabidopsis reference genome using Bowtie2, as described in ref. (Langmead and Salzberg, 2012). Genes with more than 2.0-fold change in expression and *P* value < 0.01 were considered to be differentially expressed genes. GO enrichment was determined using agriGO as described in ref. (Du et al., 2010) and Heatmap was made on https://hiplot.com.cn/.

### Quantitative RT-PCR

Total RNA was isolated from inflorescences (with mature flowers removed) using the Easy Plant RNA Kit (Easy-Do, CAT DR0406050). For DEX treatment experiments, transgenic inflorescences were treated by 10 uM DEX for 3 d. First-strand cDNA was synthesized using oligo d(T) and reverse transcriptase (Vazyme). qPCR was performed using Hieff qPCR SYBR Green master mix (Yeasen, CAT 11201ES08) in a LightCycler 480 (Roche) according to the manufacturers’ instructions. Three technical replicates were carried out. All primers used for qRT-PCR were listed in Supplemental Dataset S5. Quantification of *Actin* gene served as a control.

### ChIP assay

ChIP assays were performed as previously described in ref. (Bowler et al., 2004). The inflorescences were treated with 1% (v/v) formaldehyde to cross-link the protein-DNA complexes on ice. After isolation and sonication, samples were centrifuged at 12,000 g for 10 min at 4°C. The supernatants were precleared with 20 uL protein-A agarose beads for at least 1 hour, then incubated with 20 uL Myc-Nanoab-agarose beads (Lablead, CAT MNA-25-500) overnight. The precipitated DNA samples were quantified by qPCR using primers listed in Supplemental Dataset S5. Quantification of the *Actin* gene served as a control.

### *VGD1* expression level and pattern analysis by GUS staining

*pVGD1::GFP-GUS* was utilized to detect the expression pattern of *VGD1* in WT and *vip4-4* sepals during stage 10 to stage 14. *pVGD1::GFP-GUS* in WT and *pVGD1::GFP-GUS* in *vip4-4* were derived from the same genetic line. GUS staining was conducted following the procedure described in ref. (Sessions et al., 1999). Briefly, flowers were subjected to overnight staining with a solution containing 50 mM sodium phosphate buffer (pH 7.0), 0.2% (v/v) Triton-X-100, 10 mM potassium ferrocyanide, 10 mM potassium ferricyanide, and 1 mM X-gluc at 37 °C. Subsequently, the stained tissue was dehydrated and cleared using a series of ethanol solutions. GUS-stained sepals were then photographed using a Leica binocular.

### Image processing for growth quantification

The procedures for image processing and growth quantification were carried out following the methodology described in ref. (Hong et al., 2016; Zhu et al., 2020). In brief, the confocal stacks obtained from live imaging of sepal growth were converted from LSM to TIFF format using FIJI. The MorphoGraphX (MGX) 3D image analysis software (de Reuille et al., 2015), was utilized for growth analyses. The surfaces of the samples were detected and meshes representing the surfaces were generated. Fluorescent signals were projected onto the meshes, and the cells were segmented within the meshes.

### AFM

AFM was conducted to examine the cell wall stiffness of sepals at stage 10. The sepals were carefully removed from the flowers and placed in a solid growth medium in a Petri dish, following the protocol described in ref. (Bovio et al., 2019), with minor modifications. An R=400 nm spherical-end tip with a nominal force constant of 42 N/m was used. Prior to the measurements, each cantilever was calibrated using indentation on sapphire and thermal tune, both conducted in water.

### Osmotic treatments measuring sepal stiffness

WT and *vip4-4* sepals at stage 10 were subjected to 0.4 M NaCl treatment for 30 min. The areas of the sepals before and after the treatment were compared using MGX, following the methodology described in ref. (Hong et al., 2016; Kierzkowski et al., 2012). For IAA treatment experiments, WT inflorescences were treated by 1 mM IAA for 1 d. The flowers at stage 10 were then processed according to the above-mentioned methods.

## Acknowledgements

We thank Z. Chang for comments and suggestions on the manuscript. We thank R. Wang for drawing the schematic diagram. We thank Y. Jiao for sharing the seeds for *arf3-1*(+/-) *arf4-2*. We thank X. Liu for providing the *pARF3::ARF3-GFP*, *pARF3::ARF3m-GFP* plasmids. We thank S. Bovio and C. Lionnet from PLATIM platform (ENS de Lyon) for their help in using AFM and confocal microscope. We thank J. Li for sharing the seeds for *pPIN1::PIN1-GFP* (*Col*-0). This work was supported by National Natural Science Foundation of China grant no. 32070853 and no. 32270867, Hundred-Talent Program of Zhejiang University, China Postdoctoral Science Foundation grant no. 2021M702855, the European Research Council (ERC, grant no. 101019515, “Musix”), CEFIPRA grant no. 6103-1, the French National Research Agency through a European ERA-NET Coordinating Action in Plant Sciences (ERA-CAPS, grant No. ANR-17-CAPS-0002-01), National Institute of General Medical Sciences of the National Institutes of Health under Award Number R01GM134037. The content is solely the responsibility of the authors and does not necessarily represent the official views of the funders.

## Author contributions

Conception and design of experiments: SX, AHKR, OH, and LH. Isolation and characterization of *vip4-4* and *vip4-5* mutants: SX, and LH. BSA-seq analysis: LH and ZM. Live imaging and analysis: SX, D-CT, XH, and LH. AFM analysis: SX. RNA-seq analysis: XW, DQ and MZ, qRT-PCR analysis: SX, XH and XZ. Sepal phenotypes quantification: SX, DX and DQ. ChIP-qPCR analysis: SX and XH. Manuscript writing: SX and LH. Manuscript revising and editing: SX, XH, D-CT, XZ, XW, DQ, MZ, DX, AHKR, OH, and LH.

